# Parabens Promote Pro-Tumorigenic Effects in Luminal Breast Cancer Cell Lines with Diverse Genetic Ancestry

**DOI:** 10.1101/2022.12.01.518556

**Authors:** Jazma L. Tapia, Jillian C. McDonough, Emily L. Cauble, Cesar G. Gonzalez, Dede K. Teteh, Lindsey S. Treviño

## Abstract

One in 8 women will develop breast cancer in their lifetime. Yet, the burden of disease is greater in Black women. Black women have a 40% higher mortality rate compared to White women, and a higher incidence of breast cancer at age 40 and younger. While the underlying cause of this disparity is multifactorial, exposure to endocrine disrupting chemicals (EDCs) in hair and other personal care products has been associated with an increased risk of breast cancer. Parabens are known EDCs that are commonly used as preservatives in hair and other personal care products, and Black women are disproportionately exposed to products containing EDCs. Studies have shown that parabens impact breast cancer cell proliferation, death, migration/invasion, and metabolism, as well as gene expression *in vitro*. However, these studies were conducted using cell lines of European ancestry; to date, no studies have utilized breast cancer cell lines of West African ancestry to examine the effects of parabens on breast cancer progression. Like breast cancer cell lines with European ancestry, we hypothesize that parabens promote pro-tumorigenic effects in breast cancer cell lines of West African ancestry. Luminal breast cancer cell lines with West African ancestry (HCC1500) and European ancestry (MCF-7) were treated with biologically relevant doses of methylparaben, propylparaben, and butylparaben. Following treatment, estrogen receptor target gene expression and cell viability were examined. We observed altered estrogen receptor target gene expression and cell viability that was paraben- and cell-line specific. This study provides greater insight into the tumorigenic role of parabens in the progression of breast cancer in Black women.

## Introduction

Breast cancer is the 2^nd^ leading cause of cancer death in women [1]. Although incidence rates for both Black and White women have converged, Black women are more likely to be diagnosed with breast cancer under the age of 40, and are 40% more likely to die from the disease than White women [2, 3]. The factors underlying breast cancer disparities are multifactorial, including the significant role of sociocultural practices, such as the use of hair and other personal care products containing endocrine disrupting chemicals (EDCs) [4]. EDCs are toxins found in the environment, hair and other personal care products, and food sources. EDCs can disrupt the body’s homeostasis through interference with the endocrine system which is responsible for regulation of various biological functions, such as growth and development [5]. EDCs are thought to function by activating and disrupting estrogen receptor (ER) signaling and leading to sustained and aberrant hormone receptor activity [6]. Studies have found associations between exposure to EDCs and increased risk of breast cancer [6-9].

Parabens are a class of EDCs that can mimic estrogen in the body and are used as preservatives in hair and other personal care products. The Environmental Working Group (EWG) developed a hazardous chemical scale for beauty, personal care, and household product ingredients [10]. Using a scale of 1 (best) to 10 (worst), consumers can determine the hazard level of their products. According to the EWG, products containing butylparaben (BP) and propylparaben (PP) are scored high on the hazardous scale [6], while products containing methylparaben (MP) rank moderately [3, 10]. Hormonally hazardous chemicals have been linked to breast cancer; specifically, studies have found estrogenic properties in paraben-laden hair and other personal care products that are heavily marketed to Black women [11-13]. Parabens have been measured in biological samples from the NHANES biomonitoring study; across studies, the median/mean of PP levels are 4-15 µ g/L [14-20], the range of BP levels are 0.2-1,240 µ g/L [15, 20], and the mean of MP levels are 38-63 µ g/L [15, 17, 20-22]. There are racial/ethnic disparities in exposure to parabens. For example, 5x higher levels of MP, and 3.6x higher levels of PP were measured in Black women compared to White women [15]. These disparities have been shown to persist into more recent NHANES cycles and are most apparent in children [19]. Although there may be many factors underlying the racial disparity in paraben levels, one potential factor is prolonged use of hair and other personal care products that contain high concentrations of parabens [4, 23, 24].

The use of products containing estrogens has been linked to adverse health effects, such as premature sexual development [11]. Estrogen/estrogen receptor (ER) signaling play a role in breast cancer progression [25]. Estrogenic activity has been detected in personal care products (e.g., oil hair lotion, intensive skin lotion) commonly used by Black women, thereby potentially increasing the risk for breast cancer in this under-resourced community [11]. Studies have shown that parabens are present in normal and in tumor breast tissue [26-29]. However, these studies did not include breast tissue from Black women. Furthermore, to the best of our knowledge, studies examining the pro-tumorigenic effects of parabens *in vitro* have been conducted solely in breast cancer cell lines of European ancestry [30-37]. The lack of studies examining the effects of parabens on breast cancer progression using samples and cell lines from Black women represents a significant gap in knowledge, given that racial disparities in breast cancer risk and mortality exist, and that Black women are disproportionately exposed to hair and other personal care products containing high levels of parabens. We hypothesized that, similar to what has been shown for breast cancer cell lines with European ancestry, parabens promote pro-tumorigenic effects in breast cancer cell lines with West African ancestry. Since parabens are thought to act through ER signaling, we focused on ER+ luminal breast cancer cell lines in this study.

## Materials and Methods

### Cell Viability

Cell viability of cell lines with European ancestry MCF-7 (ATCC, CAT: HTB-22) and West African ancestry HCC1500 (ATCC, CAT: CRL-2329) was assessed utilizing CellTiter-Glo 2.0™ Luminescent Cell Viability Assay (Promega™, CAT: G9242). West African ancestry of the HCC1500 cell line was determined using established Ancestry Informative Markers [38]. Approximately 3,000 MCF-7 cells and 10,000 HCC1500 cells were seeded in black 96-well optical bottom plates (Thermo Scientific™, CAT: 165305) in 100 µL per well, using phenol red-free RPMI medium (Gibco™, CAT: 11835030) supplemented with 10% FBS (Atlanta Biologicals Inc., CAT: S11150) and incubated at 37°C. Following overnight incubation and cell adherence, the seeding medium was replaced with fresh phenol red-free RPMI medium supplemented with 10% charcoal-stripped FBS (Atlanta Biologicals Inc., CAT: S11650), and cells were incubated at 37°C for 48 hours to deplete steroid hormone levels in the cells. On day four, cells were treated with various concentrations of propylparaben (Propyl 4-hydroxybenzoate, PP; Acros Organics, CAT: 131591000), methylparaben (Methyl 4-hydroxybenzoate, MP; Acros Organics, CAT: 126961000), and butylparaben (Butyl 4-hydroxybenzoate, BP; Acros Organics, CAT: 403571000). Chemicals were dissolved in ethanol (EtOH) (Acros Organics, CAT: 615090040), diluted with phenol red-free RMPI supplemented with 10% charcoal-stripped FBS, and added to each well. Cells were incubated at 37°C for 96 hours. On day 8, the medium was replaced with fresh-treated medium. Cells were incubated at 37°C for 72 hours. On day 11, cell viability was measured using CellTiter-Glo 2.0™ Luminescent Cell Viability Assay to measure ATP, as an indicator of metabolically active cells, in either the Tecan Infinite M200 Pro or the Biotek Synergy Lx plate reader. EtOH served as a negative control, and β-Estradiol (E2; Acros Organics, CAT: 436320050) served as a positive control. The E2 concentrations used were: 0.01 nM, 0.1 nM, 1nM, 10 nM, and 100 nM. The following concentrations were used for PP, MP, and BP: 0.002 µM, 0.02 µM, 0.2 µM, 2 µM, and 20 µM. These concentrations fall within the range of what has been measured in biological samples in the NHANES biomonitoring study [14-20] and in breast tissue [21, 22, 26-29, 39]. Treatments were done in triplicates, and experiments were repeated six times.

MCF-7 and HCC1500 cells were also treated with either E2 (0.01 nM, 0.1 nM, 1nM, 10 nM, and 100 nM) or PP, MP, BP (0.002 µM, 0.02 µM, 0.2 µM, 2 µM) individually or in a mixture of all three. EtOH was used as a negative control.

### Cell Viability Assay with Estrogen Receptor Antagonist

Cell viability was measured as described above. MCF-7 and HCC1500 cells were treated with either E2 (0.1 nM, 1 nM, or 10 nM), or PP, MP, BP (0.002 µM, 0.02 µM, 0.2 µM, or 2 µM), individually with or without the ER antagonist 1 nM Fulvestrant (ICI 182,780; Tocris Bioscience, CAT: 10-471-0) to determine if the effect of parabens on cell viability is ER-mediated. EtOH was used as a negative control. Experiments were repeated four times.

### Estrogen Receptor Target Gene Expression

Cells were seeded in CytoOne 6-well plates (USA Scientific, CAT: CC7682-7506) at different densities with phenol red-free RPMI medium, supplemented with 10% FBS, and incubated at 37°C. Cell densities plated per well per cell line are as follows: MCF-7 at 3 × 10^5^ cells/well, HCC1500 at 5 × 10^5^ cells/well, BT-474 (ATCC, CAT: HTB-20) at 4 × 10^5^ cells/well, and MDA-MB-175-VII (ATCC, CAT: HTB-25) at 5 × 10^5^ cells well. BT-474 (European ancestry) and MDA-MB-VII (West African ancestry). After 24 hours, the medium was replenished to contain fresh phenol red-free RPMI medium supplemented with 10% charcoal-stripped FBS. The cells were incubated at 37°C for 48 hours. On day four, the cells were treated with charcoal-stripped phenol red-free RMPI supplemented with the appropriate concentrations of PP, MP, and BP (0.002 µM, 0.02 µM, 0.2 µM, 2 µM, and 20 µM), EtOH, or E2 (10 nM) controls, and incubated at 37°C. After 6 hours, cells were washed with 1mL PBS (Gibco™, CAT: 20012050), and 500 µL pre-warmed 0.25% Trypsin-EDTA (Gibco™, CAT: 15400054) solution was added. Following incubation and detachment from the plate, 500 mL of phenol red-free charcoal-stripped medium was added. Medium containing the detached cells was collected in 1.5 mL microcentrifuge tubes and centrifuged at 300 x G for 5 minutes. The cell pellets were stored at -80°C. EtOH served as a negative control and E2 (10 nM) served as a positive control.

Total RNA was isolated from each cell line using the RNeasy Mini Kit (Qiagen, CAT: 74106). RNA concentrations were measured using a NanoDrop One (Thermo Fisher, CAT: AZY1810576). 500 ng of RNA was reverse transcribed to cDNA by using a High-Capacity RNA-to-cDNA Kit (Applied Biosystems, CAT: 4388950). The PowerUp SYBR Green Master Mix (Applied Biosystems, CAT: A25777), in combination with primers *hGAPDH, hTFF1, hPGR, hGREB1, hMYC*, and *hCCND1* **(Table 1)**, was used for gene expression analysis of known estrogen receptor target genes using a QuantStudio™ 3 Real-Time PCR System (Applied Biosystems™, CAT: A28567). All primers were purchased from Integrated DNA Technologies. Experiments were repeated three times. The housekeeping gene, *hGAPDH* was used for normalization. The delta-delta Ct method was used to analyze the relative changes in gene expression.

**Table 1:**
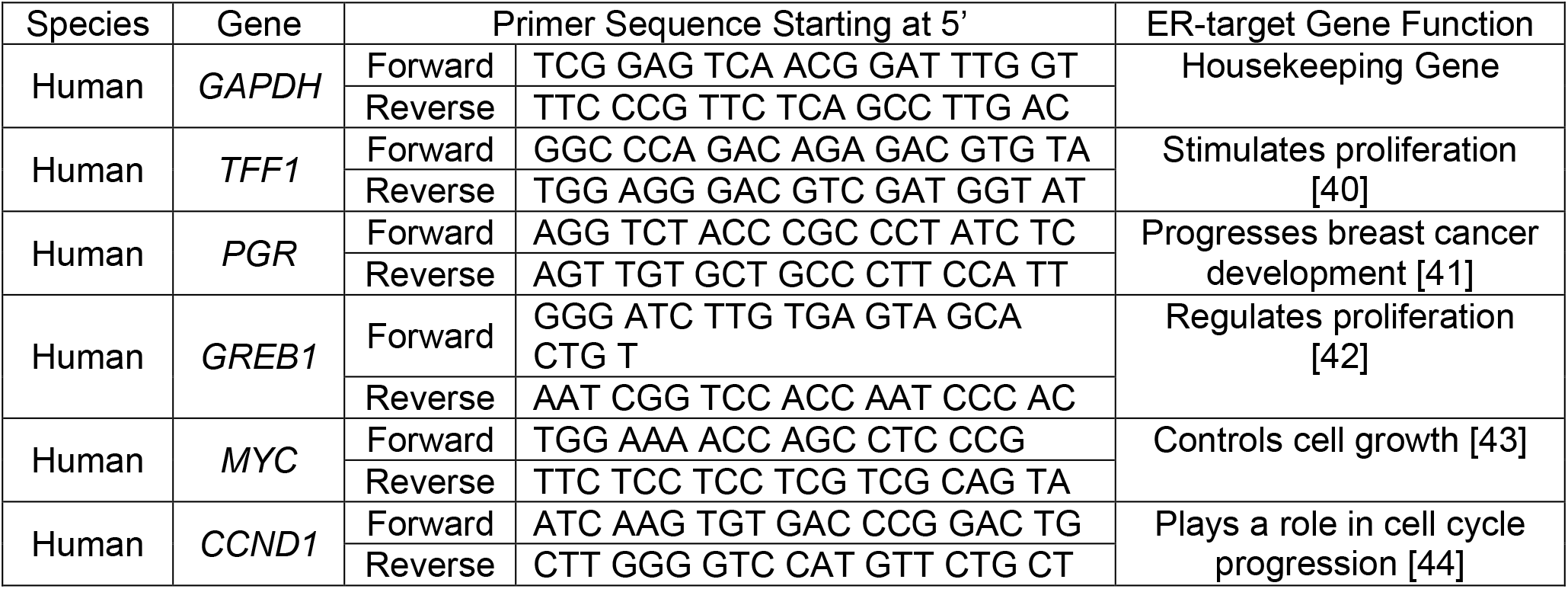
Oligonucleotide primer sequences for real-time quantitative PCR.

### Estrogen Receptor Target Gene Expression with Estrogen Receptor Antagonist

Gene expression was measured as described above. The following concentrations of PP, MP, and BP were used with and without 1 nM ICI 182,780, 2 µM, and 20 µM. EtOH, and 10 nM E2 concentrations were used with and without 1 nM ICI 182,780. Experiments were repeated four times.

### Time-Dependent Estrogen Receptor Target Gene Expression

Gene expression was measured as previously described. MCF-7 and HCC1500 cells were treated with either PP, MP, or BP (2 µM) for 1, 6, and 24 hours. EtOH was used as a negative control and E2 (10 nM) as a positive control. Experiments were repeated three times.

### Western Blot

MCF-7 and HCC1500 were seeded in 10 cm dishes (USA Scientific Inc., CAT: CC76823394) at the following densities: 2×10^6^ and 3×10^6^, respectively. All cells were incubated at 37°C. After 24 hours of incubation, the seeding medium was replaced with fresh medium containing the addition of 10% charcoal-stripped FBS. On day four, cells were treated with BP (0.002 µM, 0.02 µM, 0.2 µM, 2 µM, and 20 µM), EtOH, or E2 (10 nM) controls. Cells were harvested using cell scrapers (Fisherbrand™, CAT: 08100241), centrifuged at 120 rcf for 5 minutes, and stored in -80°C until they were used for protein extraction and western blot analysis.

Total protein was extracted from the cell lines and tissues using NP40 cell lysis buffer (Invitrogen™, CAT: FNN0021RIPA) with PMSF (Acros Organics, CAT: 215740010), a Pierce Protease Inhibitor Mini Tablet (Thermo Scientific™, CAT: A32955), and a Pierce Phosphatase Inhibitor Mini Tablet (Thermo Scientific™, CAT: A32957). Protein concentrations in the lysates were measured with the Pierce BCA Protein Assay Kit (Thermo Scientific™, CAT: 23227). Lysate mixed (10-25 μg) with 4x Sample Loading Buffer (Licor, CAT: 928-40004) and Novex™ 10X Bolt™ Sample Reducing Agent (Invitrogen™, CAT: B0009) were loaded per lane. The proteins in the lysates were separated by gel electrophoresis with Bolt™ 4-12% Bis-Tris Plus Gels (Invitrogen™, CAT: NW04122BOX) and transferred to Immobilon-FL PVDF Membranes (MilliporeSigma™, CAT: IPFL07810). Membranes were dried overnight, at room temperature, to maximize protein retention. REVERT™ Total Protein Stain (LI-COR, CAT: 926-11011) was used as a protein loading control. To block nonspecific binding, membranes were incubated with Intercept (TBS) Blocking Buffer (LI-COR, CAT: 92760001) for 1 hour at room temperature, then incubated overnight at 4 °C with the following antibody: ERα (Dilution 1:1 µg/mL; Invitrogen, CAT: MA5-13191). IRDye secondary antibody (Dilution 1:20,000; LI-COR, CAT: 82708364, 92968171, 92632214) was used to visualize the target protein with an Odyssey Classic (LI-COR). Experiments were repeated six times.

### Statistical Analysis

GraphPad Prism Software (Dotmatics, Version 8) was used to determine statistical significance. One-way ANOVA was performed for cell viability and gene expression assays. Two-way ANOVA was performed for cell viability and gene expression assays that included co-treatment with ICI 182,708. Differences were considered statistically significant when the p-value was less than or equal to 0.05. Error bars represent standard deviation (SD).

## Results

### Estrogen receptor (ER)-target genes are regulated by parabens in a dose-dependent manner

To test the effects of paraben treatment on ER-target gene expression, MCF-7 (European ancestry) and HCC1500 (West African ancestry) cells were treated with MP, PP, or BP at the doses indicated for 6 hours. Expression of well-known ER-target genes trefoil factor-1 (*TFF1*), progesterone receptor (*PGR*), growth regulating estrogen receptor binding 1 (*GREB1*), *MYC* protooncogene (*MYC)*, and cyclin D1 (*CCND1*) was measured via real-time quantitative polymerase chain reaction (RT-qPCR). Treatment with 2 µM PP significantly increased *TFF1* gene expression in HCC1500, but not MCF-7 cells **(Figure 1A)**. Treatment with 20 µM PP significantly increased *PGR* gene expression in HCC1500, but not MCF-7 cells **(Figure 1B)**. Significantly increased expression of *TFF1* was observed with 2 µM BP treatment in both MCF-7 and HCC1500 cells **(Figure 1C)**. Treatment with 2 and 20 µM BP significantly increased *PGR* gene expression in MCF-7 cells, while treatment with 20 µM BP significantly increased *PGR* gene expression in HCC1500 cells **(Figure 1D)**. Paraben- and cell-line dependent results were also observed for *GREB1* **(Supplemental Figure 1A and D)**, *MYC* (**Supplemental Figure 1B and E)**, and *CCND1* **(Supplemental Figure 1C and F)**. Treatment with MP significantly increased *PGR* (20 µM) and *CCND1* (0.002, 0.02, 0.2, and 2 µM) compared to control in HCC1500, but not MCF-7, cells (**Supplemental Figure 2)**. These results suggest that PP and BP may be more estrogenic than MP in luminal breast cancer cells.

**Figure 1.**
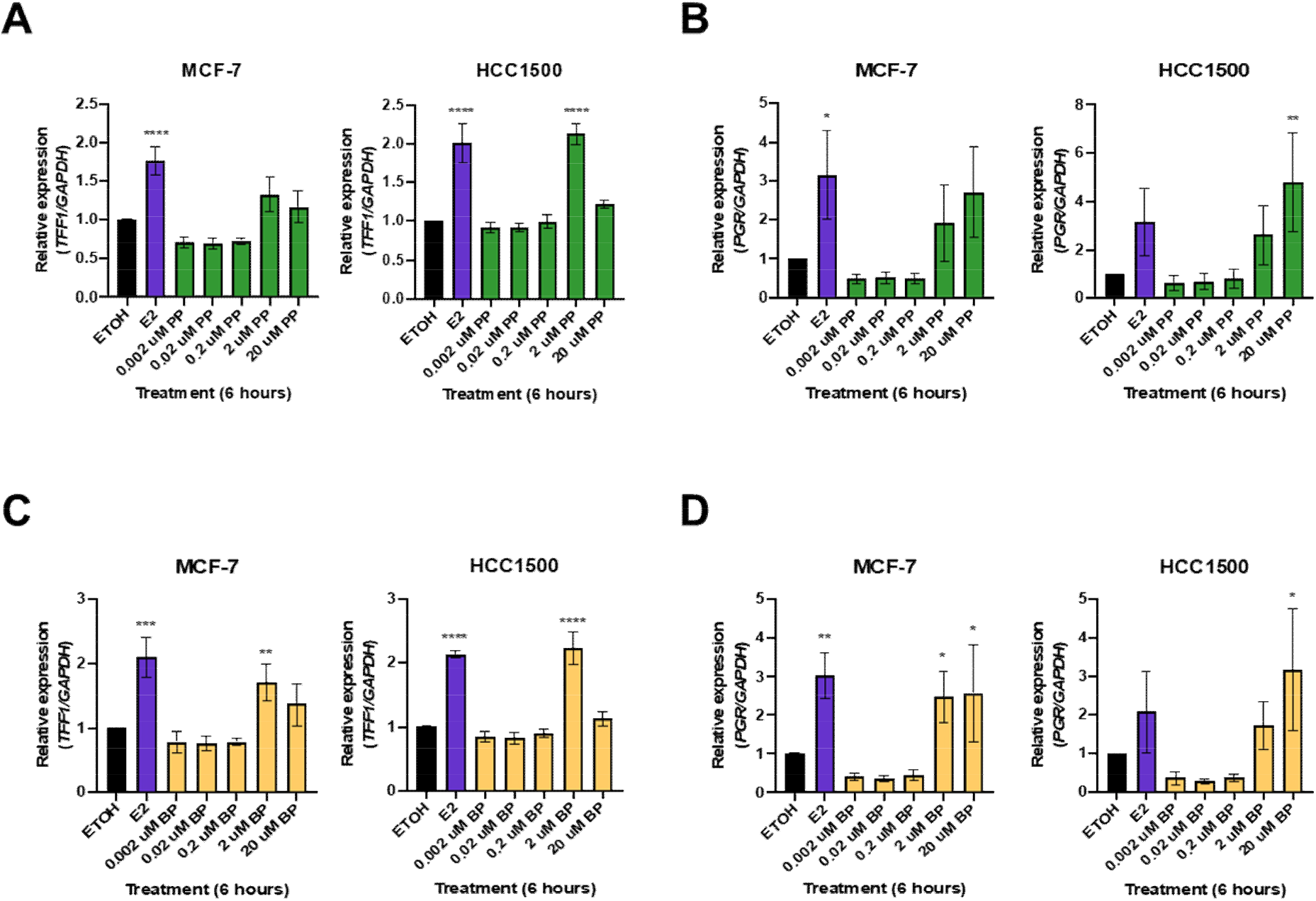
PP- and BP-mediated regulation of ER target gene expression in MCF-7 and HCC1500. MCF-7 and HCC1500 luminal A breast cancer cell lines were treated with the indicated doses of PP or BP for 6 hours. **(A-D)** Quantification of **(A, C)** *TFF1* and **(B, D)** *PGR* gene expression following PP or BP exposure, respectively. Ethanol (EtOH) was used as a negative control, and estradiol (E2, 10 nM) served as a positive control. GAPDH was used as a housekeeping gene. n=3, *p<0.05, **p<0.01, ***p<0.001, ****p<0.0001, one-way ANOVA.

To determine whether the estrogenic activity of PP and BP was cell-line specific, gene expression was measured in additional luminal cell lines, BT-474 (European ancestry), and MDA-MB-175-VII (West African ancestry), following paraben exposure. Treatment with PP or BP did not increase *TFF1* gene expression in either cell line **(Supplemental Figure 3A and 4A**). Treatment with 20 µM PP or BP significantly increased *GREB1* gene expression in BT-474, but not MDA-MB-175-VII, cells **(Supplemental Figure 3B and 4B**). Treatment with 20 µM PP or 20 µM BP significantly increased *MYC* expression in both cell lines **(Supplemental Figure 3C and 4C**). Treatment with 2 µM BP also significantly increased *MYC* gene expression in BT-474 cells **(Supplemental Figure 4C**). Additionally, treatment with PP, in both cell lines, significantly increased *CCND1* gene expression at all doses except 20 µM **(Supplemental Figure 3D)**. Furthermore, BP treatment significantly increased *CCND1* gene expression in both cell lines, except at the 20 µM concentration in the BT-474 cell line **(Supplemental Figure 4D)**. Taken together, these results suggest that parabens regulate ER target gene expression in a variety of luminal breast cancer cell lines.

### Paraben-mediated effects on ER target gene expression are ER-dependent

To determine whether the observed PP- and BP-mediated effects on ER-target gene expression are ER-dependent, MCF-7 and HCC1500 cells were co-treated with PP or BP and ICI 182,780 (Fulvestrant) for 6 hours. Co-treatment with ICI 182,780 blocked the PP-mediated increase in *TFF1, PGR, MYC*, and *CCND1* expression in both cell lines and in *GREB1* expression in HCC1500 cells **(Figure 2)**. Co-treatment with ICI 182,780 blocked the BP-mediated increase in *TFF1, PGR, GREB1, MYC*, and *CCND1* in both cell lines **(Figure 3)**. It should be noted that 1 nM ICI 182,780 is not enough to block the increased ER-target gene expression observed with 10 nM E2 treatment but is sufficient to block paraben-mediated ER-target gene expression. Collectively, the results suggest that regulation of ER target gene expression by PP and BP is ER-dependent.

**Figure 2.**
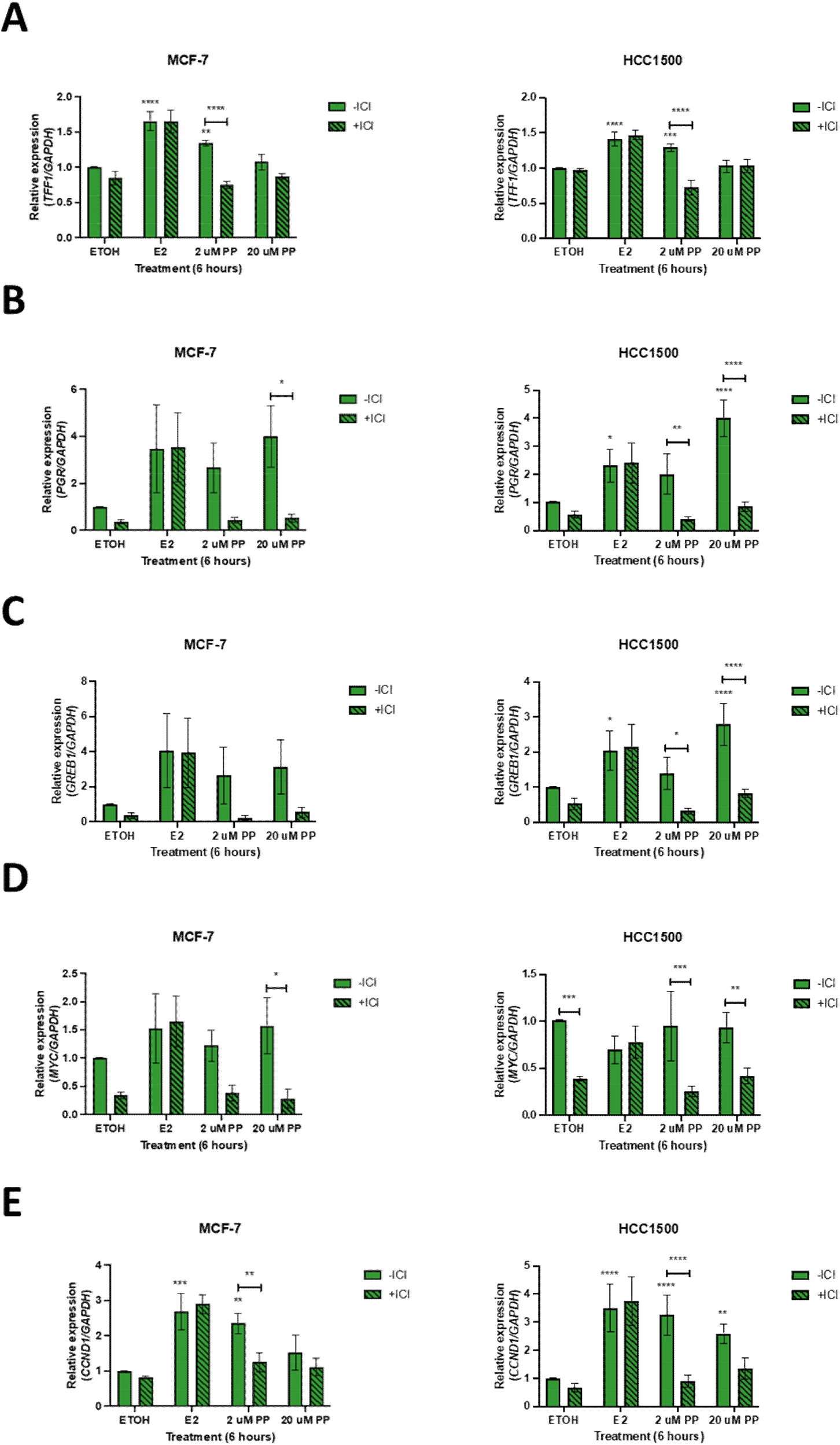
PP-mediated regulation of ER target gene expression is ER-dependent. MCF-7 and HCC1500 luminal A breast cancer cell lines were treated with 2 µM or 20 µM PP for 6 hours in the presence or absence of ER-antagonist, ICI 182,780 (1 nM). **(A-E)** Relative gene expression of **(A)** *TFF1*, **(B)** *PGR*, **(C)** *GREB1*, **(D)** *MYC*, and **(E)** *CCND1* was assessed by real-time qualitative polymerase chain reaction. EtOH was used as a negative control, and E2 (10 nM) served as a positive control. GAPDH was used as a housekeeping gene. n=4, *p<0.05, **p<0.01, ***p<0.001, ****p<0.0001, two-way ANOVA.

**Figure 3.**
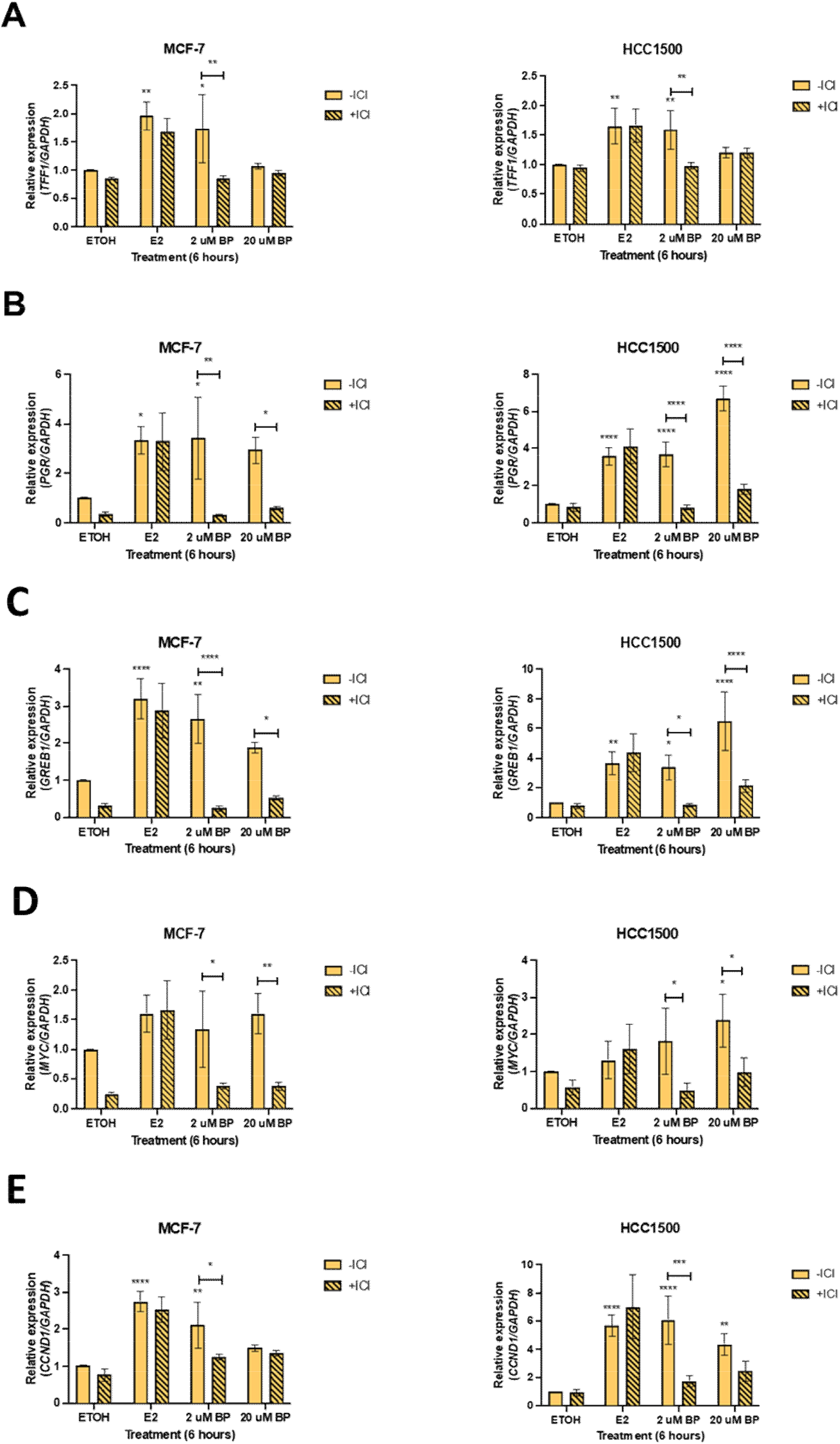
BP-mediated regulation of ER target gene expression is ER-dependent. MCF-7 and HCC1500 luminal A breast cancer cell lines were treated with 2 µM or 20 µM BP for 6 hours in the presence or absence of ER-antagonist, ICI 182, 780 (1 nM). **(A-E)** Relative gene expression of **(A)** *TFF1*, **(B)** *PGR*, **(C)** *GREB1*, **(D)** *MYC*, and **(E)** *CCND1* was assessed by real-time qualitative polymerase chain reaction. EtOH was used as a negative control, and E2 (10 nM) served as a positive control. GAPDH was used as a housekeeping gene. n=4, *p<0.05, **p<0.01, ****p<0.0001, two-way ANOVA.

### ER-target genes are regulated by PP and BP in a time-dependent manner

To examine time-dependent regulation of ER target genes by parabens, luminal breast cancer cells (i.e., MCF-7 and HCC1500) were treated with PP, BP, or MP (2 µM) for either 1, 6, or 24 hours. As expected, we observed time-dependent expression of ER-target genes in both cell lines treated with estradiol (E2, 10 nM) **(Supplemental Figure 6)**. Treatment with PP significantly increased expression of *PGR* (1 and 6 hours), *GREB1* (6 hours), and *MYC* (1 hour), but not *TFF1* or *CCND1*, in MCF-7 cells **(Figure 4A-E)**. Treatment with PP significantly increased expression of *TFF1* (6 and 24 hours), *GREB1* (6 hours), and *CCND1* (1 hour), but not *PGR*, in HCC1500 cells **(Figure 4A-C, E)**. *MYC* expression was significantly decreased upon treatment with PP at the 6- and 24-hour timepoints in HCC1500 cells **(Figure 4D)**. Expression of *TFF1* was significantly decreased with BP treatment for 1 hour in MCF-7 cells **(Figure 5A)**. Treatment with BP increased expression of *PGR* (6 and 24 hours), *GREB1* (6 hours), and *MYC* (1 hour), but not *CCND1*, in MCF-7 cells **(Figure 5B-E)**. Treatment with BP increased *TFF1* (6 and 24 hours), *PGR* (6 hours), *GREB1* (6 hours), *MYC* (1 hour), and *CCND1* (1 and 6 hours) in HCC1500 cells **(Figure 5)**. Expression of *MYC* was significantly decreased upon treatment with BP for 24 hours in HCC1500 cells **(Figure 5D)**. No significant change in *TFF1* expression was observed following treatment with MP at any timepoint, in both cell lines **(Supplemental Figure 7A)**. However, treatment with MP for 1 hour significantly increased expression of *PGR* and *CCND1* in MCF-7 cells **(Supplemental Figure 7B and E)** and *CCND1* in HCC1500 cells **(Supplemental Figure 7E)**. Expression of *MYC* was significantly decreased in MCF-7 cells treated with MP for 6 and 24 hours **(Supplemental Figure 7D)**. Expression of *PGR, GREB1*, and *MYC* was significantly decreased in HCC1500 cells treated with MP for 1, 6, and 24 hours **(Supplemental Figure 7B-D)**. Together, the data suggest that parabens alter ER target gene expression in a time-dependent manner.

**Figure 4.**
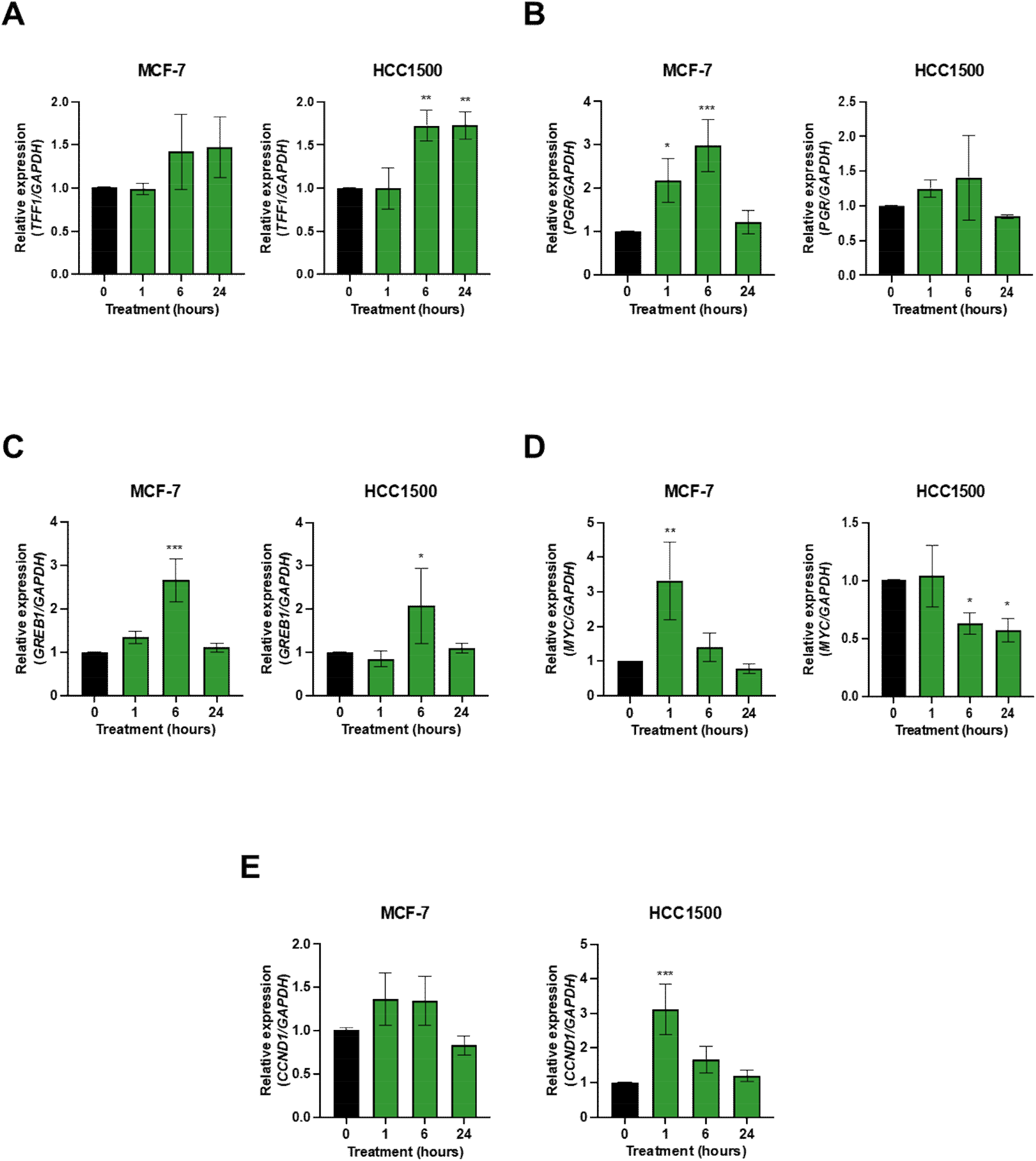
PP-mediated regulation of ER target gene expression is time dependent. MCF-7 and HCC1500 luminal A breast cancer cells were treated with 2 µM PP for 1, 6, or 24 hours. **(A-E)** Relative gene expression of **(A)** *TFF1*, **(B)** *PGR*, **(C)** *GREB1*, **(D)** *MYC*, and **(E)** *CCND1* was assessed by real-time qualitative polymerase chain reaction. EtOH was used as a negative control, and E2 (10 nM) served as a positive control. GAPDH was used as a housekeeping gene. n=3, *p<0.05, **p<0.01, ***p<0.001, one-way ANOVA.

**Figure 5.**
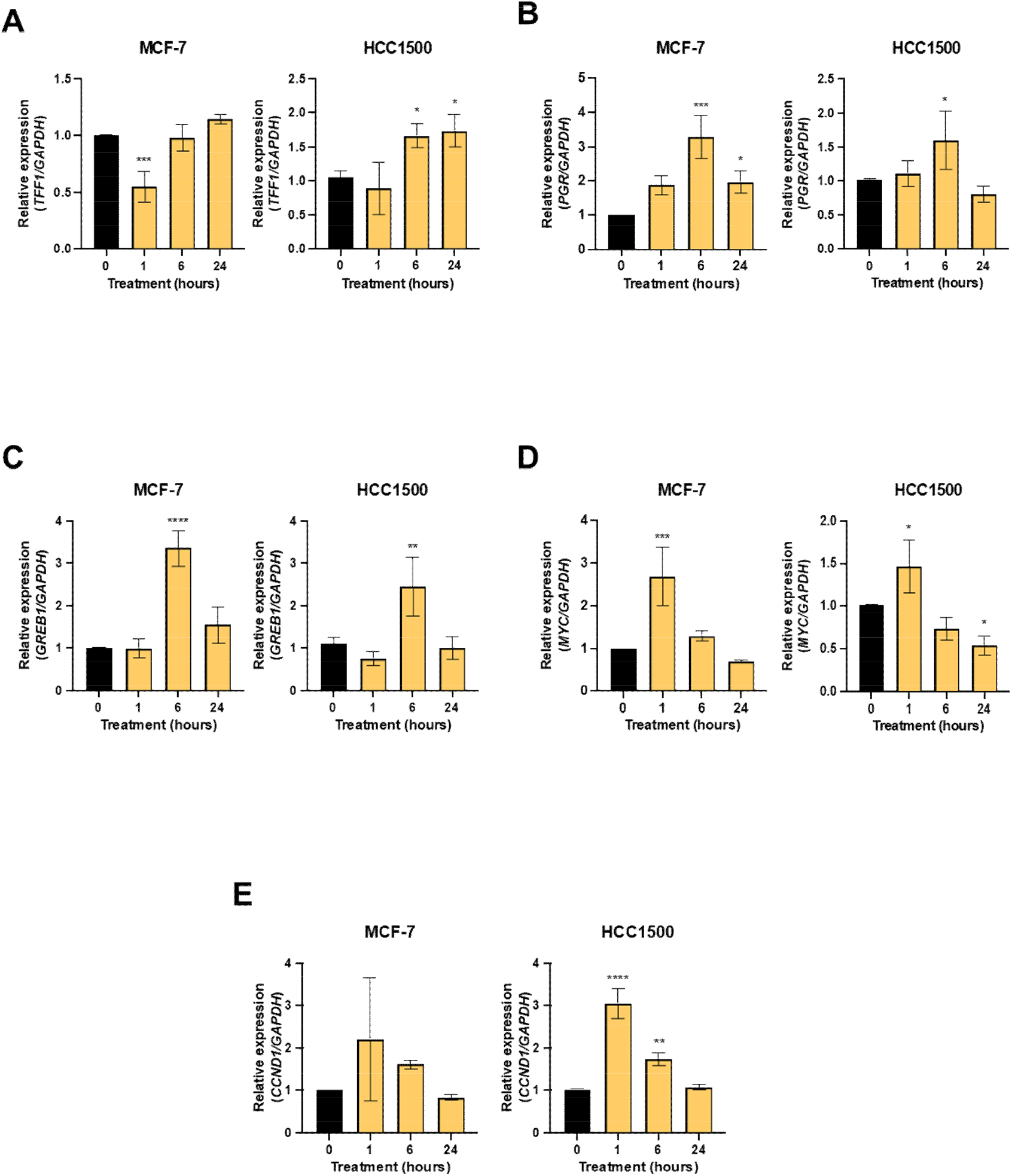
BP-mediated regulation of ER target gene expression is time dependent. MCF-7 and HCC1500 luminal A breast cancer cells were treated with 2 µM BP for 1, 6, or 24 hours. **(A-E)** Relative gene expression of **(A)** *TFF1*, **(B)** *PGR*, **(C)** *GREB1*, **(D)** *MYC*, and **(E)** *CCND1* was assessed by real-time qualitative polymerase chain reaction. EtOH was used as a negative control, and E2 (10 nM) served as a positive control. GAPDH was used as a housekeeping gene. n=3, *p<0.05, **p<0.01, ***p<0.001, ****p<0.0001, one-way ANOVA.

**Figure 6.**
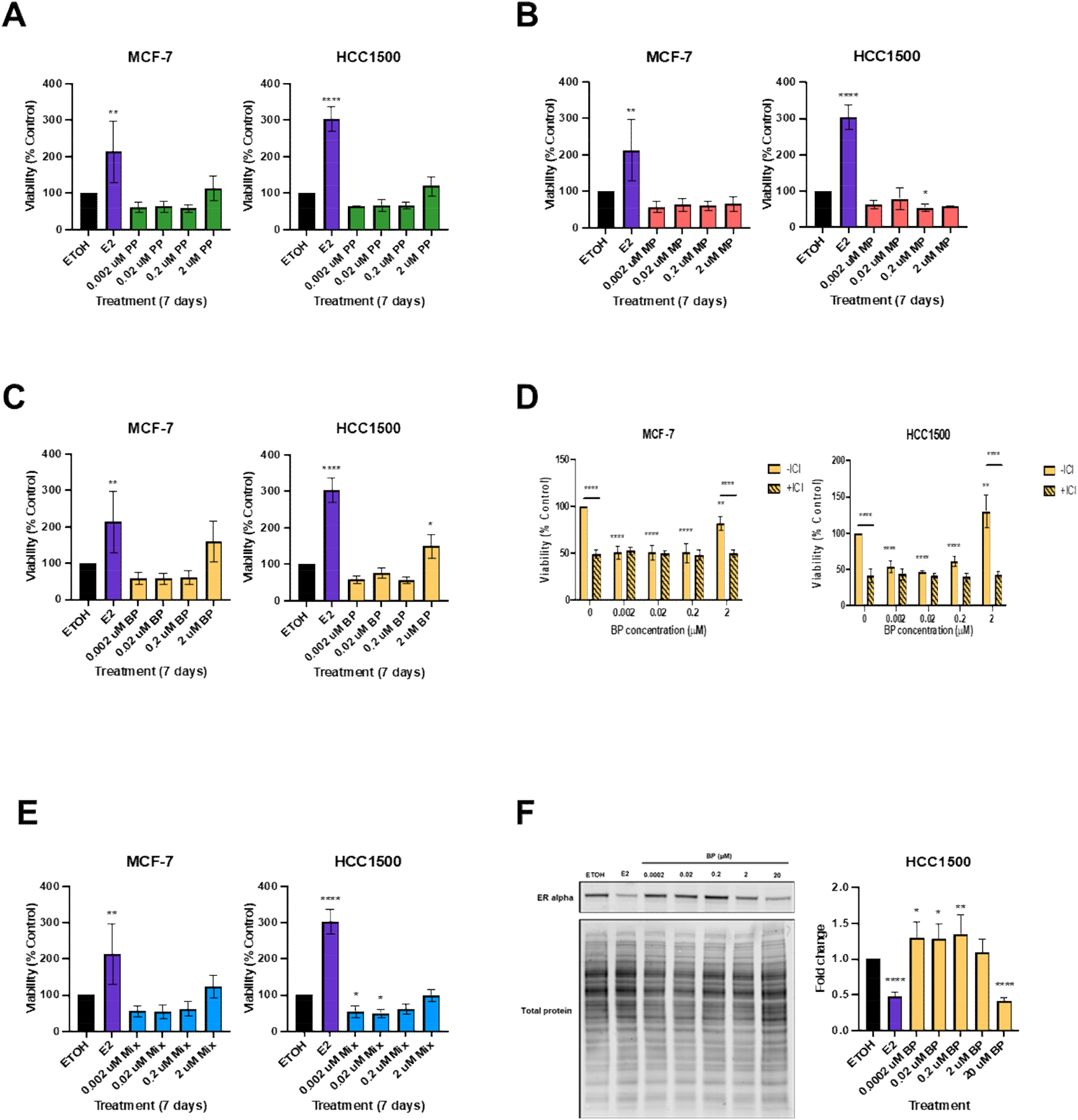
Effect of paraben exposure on cell viability is not ER-dependent. MCF-7 and HCC1500 luminal A breast cancer cells were treated with the indicated doses of **(A)** PP, **(B)** MP, or **(C)** BP for 7 days. **(D)** Effect of BP on cell viability with co-treatment with ER-antagonist, ICI 182,780 (Fulvestrant, 1 nM) for 7 days. **(E)** Effect of a mixture of PP, MP, and BP at the indicated doses for 7 days on cell viability **(F)** HCC1500 cells were treated with the indicated doses of BP for 24 hours. Representative western blot image, and quantified bar graph displaying BP-mediated effect on ERα protein expression in the HCC1500 cell line (n=6). EtOH was used as a negative control, and E2 (10 nM) serves as a positive control. n=4, *p<0.05, **p<0.01, ***p<0.001, ****p<0.0001, **A-C, E)** one-way ANOVA and **D, F)** two-way ANOVA.

### Butylparaben-mediated effects on cell viability are not ER-dependent

To test the effects of parabens on cell viability, MCF-7 and HCC1500 cells were treated with PP, BP, or MP at the indicated doses for 7 days. Treatment with either PP or MP did not increase cell viability in either MCF-7 nor HCC1500 cells **(Figure 6A, B)**. Treatment with BP (2 µM) significantly increased cell viability in the HCC1500 cell line but not in the MCF-7 cell line **(Figure 6C)**.

To investigate whether the observed effects of parabens on cell viability is mediated through ER, MCF-7 and HCC1500 cells were co-treated with either PP, BP, or MP in the presence or absence of 1 nM ER ICI 182,780. While we observed significantly decreased BP-mediated cell viability in the presence of ICI 182,780 in HCC1500 cells, decreased basal cell viability was also observed, suggesting that BP effects on cell viability are not ER-mediated **(Figure 6D)**. As expected, treatment with E2 (10 nM) significantly increased cell viability of MCF-7 and HCC1500 cells at all doses tested **(Supplemental Figure 8A)**. Like what was observed with BP-mediated cell viability in HCC1500, treatment with ICI 182,780 significantly inhibited both basal cell viability and E2-mediated cell viability **(Supplemental Figure 8A)**. Co-treatment with ICI 182,780 did not reverse the decreased cell viability observed in MCF-7 and HCC1500 cells treated with either PP or MP **(Supplemental Figure 8B and C)**.

Hair and other personal care products commonly contain more than one paraben; therefore, we determined the effect of treatment with a mixture of PP, BP, and MP for 7 days on cell viability. We observed that the BP-mediated increase in HCC1500 cell viability was blunted in the presence of PP and MP at all doses tested **(Figure 6E)**. Collectively, these data suggest that BP increases HCC1500 cell viability in an ER-independent manner.

### Butylparaben treatment stabilizes ER protein expression in luminal breast cancer cells

To determine whether butylparaben treatment regulates ER protein expression, we performed Western blot analysis. As expected, treatment with E2 (10 nM) significantly decreased ERα protein expression compared to control in the HCC1500 cell line **(Figure 6F)**. In contrast, treatment with BP (0.0002, 0.02, and 0.2 µM) significantly increased ERα protein expression compared to control in the HCC1500 cell line **(Figure 6F)**. Treatment with BP (0.0002, 0.02, 0.2, and 2 µM) also significantly increased ERα protein expression in MCF-7 cells **(Supplemental Figure 9A)**. We did not, however, observe a significant decrease in ERα protein expression as expected with E2 treatment (10 nM) in MCF-7 cells due to variability across biological replicates **(Supplemental Figure 9A)**. Nonetheless, these data suggest that BP exposure may induce prolonged activation of ERα in luminal breast cancer cells.

## Discussion

To our knowledge, this is the first study to examine the effects of paraben treatment on luminal breast cancer cells of West African ancestry *in vitro*. We know that Black women are at higher risk for developing breast cancer at younger ages, are more likely to be diagnosed with more aggressive disease and are more likely to die from breast cancer than their White counterparts [2, 45], yet previous studies examining the pro-tumorigenic effects of parabens solely focused on breast cancer cell lines of European ancestry. This lack of inclusion in breast cancer research hinders advances in prevention and therapeutic strategies that will eliminate breast cancer disparities [6]. Therefore, the goal of this study was to examine pro-tumorigenic effects of parabens in breast cancer cell lines with West African ancestry. Our data provides the foundation for further studies examining the mechanism of paraben action in Black breast cancer cells, as well as for intervention strategies for reducing exposure to hair and other personal care products that contain parabens and other harmful EDCs in Black women.

It is known that estrogens play a role in breast cancer development and progression [25]. Since parabens are thought to act as estrogens, we examined the effects of paraben treatment on ER target gene expression in diverse luminal breast cancer cell lines. ER regulates expression of its target genes in a variety of mechanisms, including through direct binding to DNA at estrogen response elements (EREs) or via tethering to other transcription factors, such as AP1 and Sp1 [46-48]. We examined the expression of genes that are regulated by direct binding to EREs (*TFF1, PGR*, and *GREB1*) or via tethering (*MYC* and *CCND1*) and found that parabens regulate ER target gene expression in a paraben-specific manner. In general, treatment with PP and BP for 6 hours exerts a greater effect on expression of ER target genes compared to MP in all cell lines tested. Interestingly, MP treatment for 6 hours increased *CCND1* expression in HCC1500 (West African ancestry), BT-474 (European ancestry), and MDA-MB-175-VII (West African ancestry) cells, suggesting that MP may be as estrogenic as PP and BP in some contexts. These results are supported by other studies that found PP and BP to be more estrogenic compared to their paraben counterparts in murine, and in breast tumors in women of European ancestry [26, 49-51]. These data also parallel the EWG hazardous chemical score for the individual parabens, where PP and BP are higher on the hazardous scale compared to MP [52-55].

Estradiol regulates gene expression with distinct time-course patterns in human breast cancer cells [56-58], so we examined time-dependent regulation of ER-target gene expression by parabens to determine whether paraben-mediated effects are time-dependent. In general, direct ER targets (*TFF1, PGR*, and *GREB1*) are induced by PP or BP with either 6 or 24 hours of treatment. Indirect ER targets (*MYC* and *CCND1*) are induced by PP or BP (and MP for *CCND1*) with 1 hour of treatment. Rapid induction of *MYC* expression by PP (MCF-7) and BP (MCF-7 and HCC1500) treatment for 1 hour, is consistent with another study that reported induction of *MYC* at ∼30 minutes following E2 exposure and regulation of expression via tethering to AP-1 [46]. The observed paraben-mediated effects on ER target gene expression are consistent with the pattern seen with E2 treatment and the respective mechanisms of regulation for either direct or indirect ER target genes. Future studies should examine the effect of parabens on ER and coregulator recruitment to direct target genes, as well as activation of cell signaling pathways and coregulator recruitment to indirect target genes.

We also observed that parabens regulate ER target gene expression in a cell-line specific manner. In general, the HCC1500 (West African ancestry) luminal A breast cancer cell line seems to be more sensitive to parabens compared to the MCF-7 (European ancestry) luminal A breast cancer cell line. We observed increased expression of *TFF1, PGR*, and *CCND1* with PP (6 hours), increased expression of *GREB1* with BP treatment (6 hours), and increased expression of *PGR* and *CCND1* with MP treatment (6 hours) in HCC1500, but not MCF-7, cells. Paraben-mediated regulation of ER target gene expression is not limited to MCF-7 and HCC1500 luminal A breast cancer cells. We also observed altered ER target gene expression in the luminal B breast cancer cell lines, BT-474 (European ancestry) and MDA-MB-175-VII (West African ancestry). In general, regulation of ER target gene expression by PP or BP was similar in BT-474 and MDA-MB-175-VII cells, except for *GREB1* and *MYC* which are not upregulated by E2 treatment (6 hours) in MDA-MB-175-VII cells. Treatment with PP or BP increased expression of *GREB1* in BT-474, but not MDA-MB-175-VII, cells. *MYC* gene expression was significantly increased in both BT-474 and MDA-MB-175-VII cell lines with PP (20 µM) or BP treatment (2 µM or 20 µM). It should be noted that *PGR* is not expressed in MDA-MB-175-VII; therefore, results for *PGR* expression in BT-474 are not shown.

Estrogens are known to regulate cell proliferation in ER+ breast cancer cells [30, 42, 59]. Studies have found that parabens interact with ER to regulate *MYC* and *CCND1* gene expression in breast cancer and non-malignant breast cells [33, 60]. *MYC* controls cell growth by promoting the cell cycle and activating cyclins, while *CCND1* plays a role in cell cycle progression [43, 44]. Parabens increase cell growth in breast cancer cell lines with European ancestry [31, 61]; therefore, we also measured the effects of parabens on cell viability in a luminal A breast cancer cell line with West African ancestry (HCC1500). We observed a paraben- and cell line-specific effect on cell viability. Specifically, BP increased cell viability in HCC1500, but not MCF-7, cells. There was no effect of PP and MP on cell viability in either cell line. Once again, this is in line with previous studies that reported BP is more estrogenic compared to the other two parabens [49, 50, 62]. Interestingly, the BP-mediated effect on cell viability in HCC1500 is not ER-mediated, suggesting that an alternative mechanism is responsible for this effect. These alternative mechanisms include, but are not limited to, 1) through interaction with the membrane-bound G-protein coupled estrogen receptor, 2) crosstalk with growth factor signaling pathways such as human epidermal growth factor receptor 2, or 3) by inducing aromatase through regulation of local estrogen levels [60, 62-64]. For example, Pan *et al* found that breast cancer cell (BT-474: European ancestry) proliferation was stimulated when BP and heregulin were combined [34], Whether these alternative mechanisms are responsible for the BP-mediated effects on cell viability in breast cancer cell lines of West African ancestry remains to be further investigated.

We utilized luminal A (MCF-7 and HCC1500) and luminal B (BT-474 and MDA-MB-175-VII) breast cancer cell lines for these experiments. The diversity of cell lines in this study was limited by the number of commercially available breast cancer cell lines of each tumor subtype with confirmed West African ancestry [65]. This limited resource presents a challenge when examining genes and pathways involved in adverse EDC exposures in the context of health disparities [38, 65]. Therefore, examining the ancestry of commonly used cell lines is warranted to adequately model the disparity of breast cancer risk and outcomes within laboratory studies [6]. Prior to this study, West African ancestry of the HCC1500 cell line was confirmed using established Ancestry Informative Markers [38]. Another limitation of this study is the lack of a detailed mechanism underlying the observed effects on ER target gene expression and cell viability due to paraben treatment. We observed that paraben-mediated effects on ER target gene expression, but not cell viability, are ER-dependent. We also observed that BP stabilizes ER protein expression in luminal A breast cancer cell lines, suggesting prolonged and abnormal activation of ER upon exposure. However, the exact mechanisms of paraben action are beyond the scope of this paper and will be explored in future studies.

## Conclusion

In this study, we observed paraben-mediated changes in ER target gene expression and cell viability that are comparable to those changes observed with estradiol treatment. Given that Black women exhibit higher burden of parabens, our results demonstrating pro-tumorigenic effects of parabens in luminal breast cancer cell lines of West African ancestry have translational relevance. Our data highlight the need for culturally relevant intervention strategies to reduce adverse exposure to parabens and other EDCs in hair and other personal care products, thus addressing and reducing known breast cancer disparities in Black women at higher risk for developing the disease.

## Acknowledgments

The current laboratory study is a part of the *Bench to Community Initiative* [66] which was initiated in response to community stakeholder questions about the biological effects of chemicals found in commonly used products by Black women. BCI is composed of multidisciplinary collaborators bringing together basic researchers in endocrinology (Treviño), social-behavioral science (Teteh), and community stakeholders (D. Bing Turner, MPH, Maggie Hawkins, MPH, CHES, Tiah Tomlin-Harris, MS, and Tonya Fairley, Licensed Trichologist). Bringing together this multidisciplinary research collaborative provides unique perspectives to approach breast cancer disparities while reducing adverse EDC exposure in hair and other personal care products used by Black women.

## Figure Legends

**Supplemental Figure 1.**
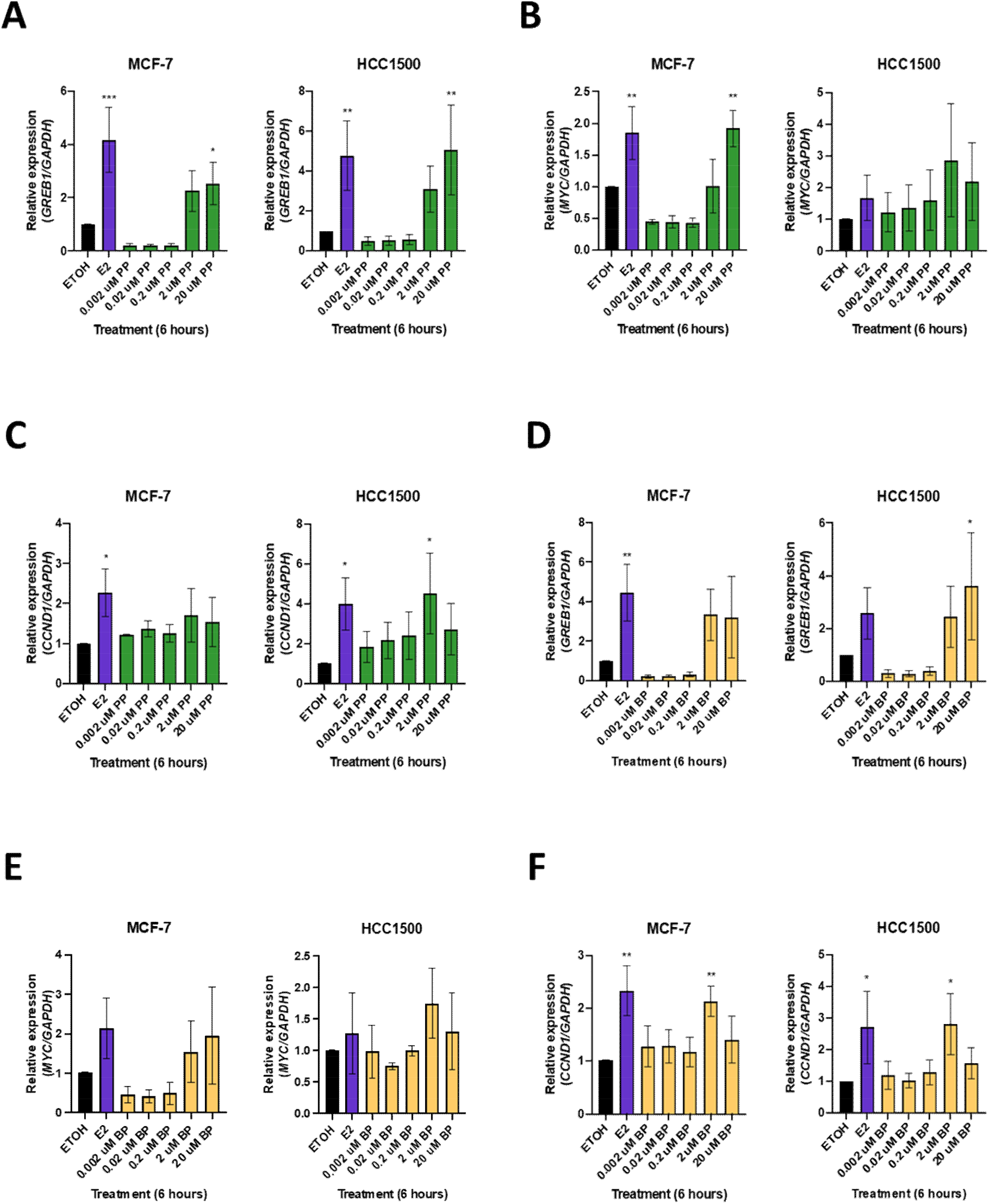
PP- and BP-mediated regulation of ER target gene expression in MCF-7 and HCC1500 cells. MCF-7 and HCC1500 luminal A breast cancer cells were treated with the indicated doses of PP or BP for 6 hours. Quantification of **(A, D)** *GREB1*, **(B, E)** *MYC*, **and (C, F)** *CCND1* gene expression following PP or BP exposure, respectively. EtOH was used as a negative control, and E2 (10 nM) served as a positive control. GAPDH was used as a housekeeping gene. n=3, *p<0.05, **p<0.01, ***p<0.001, one-way ANOVA.

**Supplemental Figure 2.**
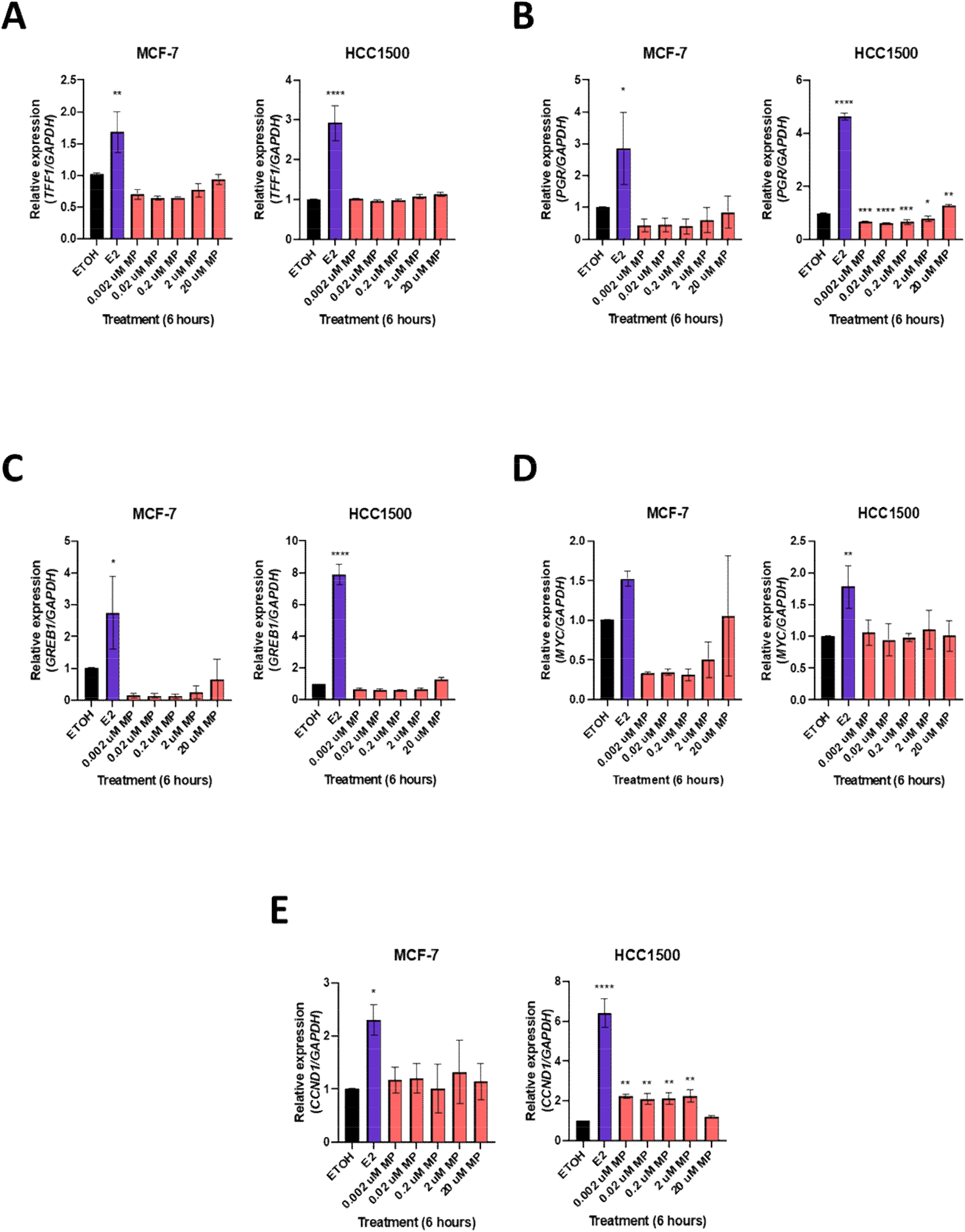
MP has minimal to no effect on expression of ER target genes in MCF-7 and HCC1500 cells. MCF-7 and HCC1500 luminal A breast cancer cells were treated with the indicated doses of MP for 6 hours. Quantification of **(A)** *TFF1*, **(B)** *PGR*, **(C)** *GREB1*, **(D)** *MYC*, and **(E)** *CCND1* gene expression following MP exposure. EtOH was used as a negative control, and E2 (10 nM) served as a positive control. GAPDH was used as a housekeeping gene. n=3, *p<0.05, **p<0.01, ***p<0.001, one-way ANOVA.

**Supplemental Figure 3.**
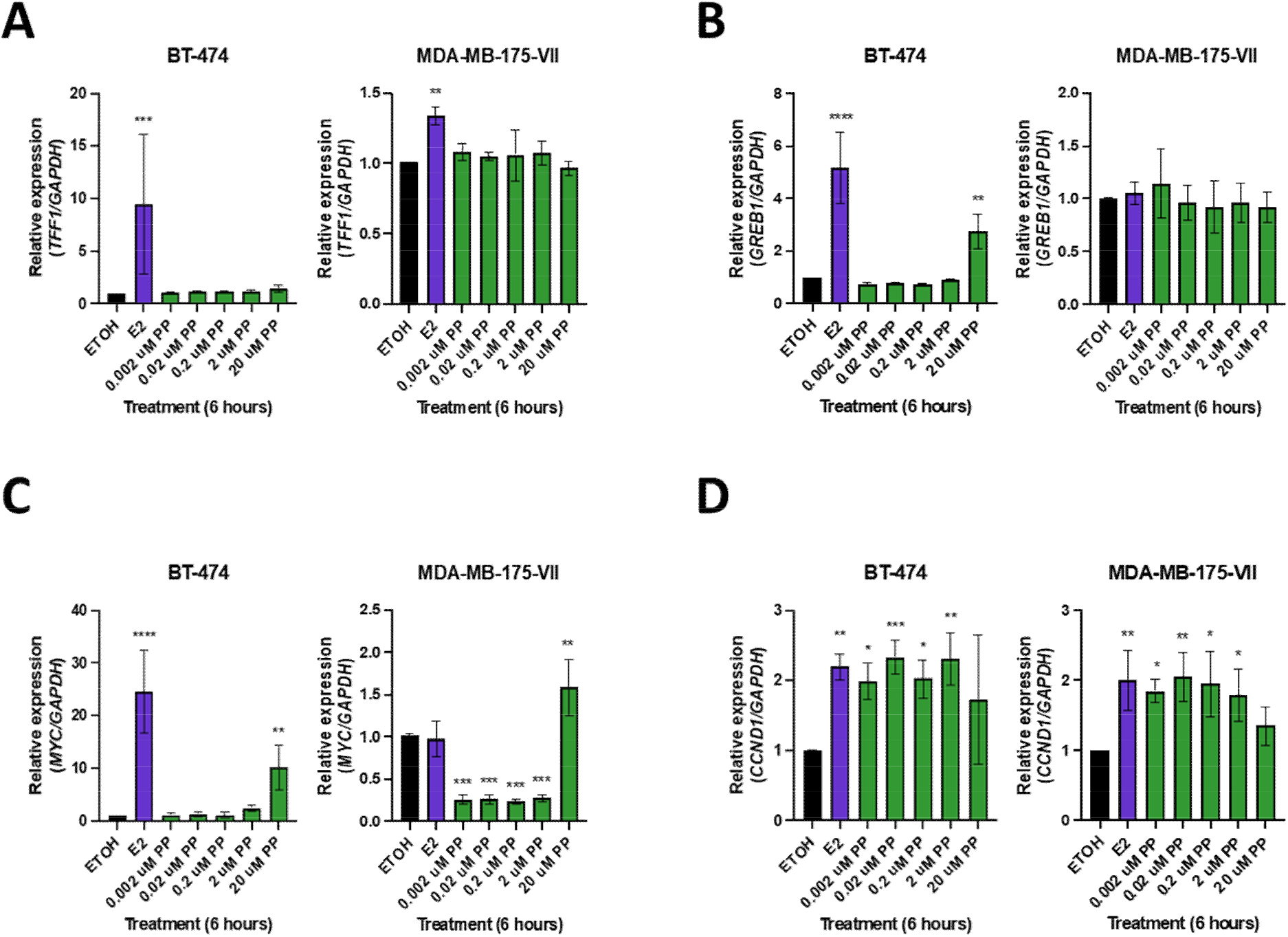
PP-mediated regulation of ER target gene expression in BT-474 and MDA-MB-175-VII cells. BT-474 and MDA-MB-175-VII luminal B breast cancer cells were treated with the indicated doses of PP for 6 hours. Quantification of **(A)** *TFF1*, **(B)** *GREB1*, **(C)** *MYC*, and **(D)** *CCND1* gene expression following PP exposure. *PGR* was not expressed in the MDA-MB-175-VII cell line. EtOH was used as a negative control, and E2 (10 nM) served as a positive control. GAPDH was used as a housekeeping gene. n=3, *p<0.05, **p<0.01, ***p<0.001, one-way ANOVA.

**Supplemental Figure 4.**
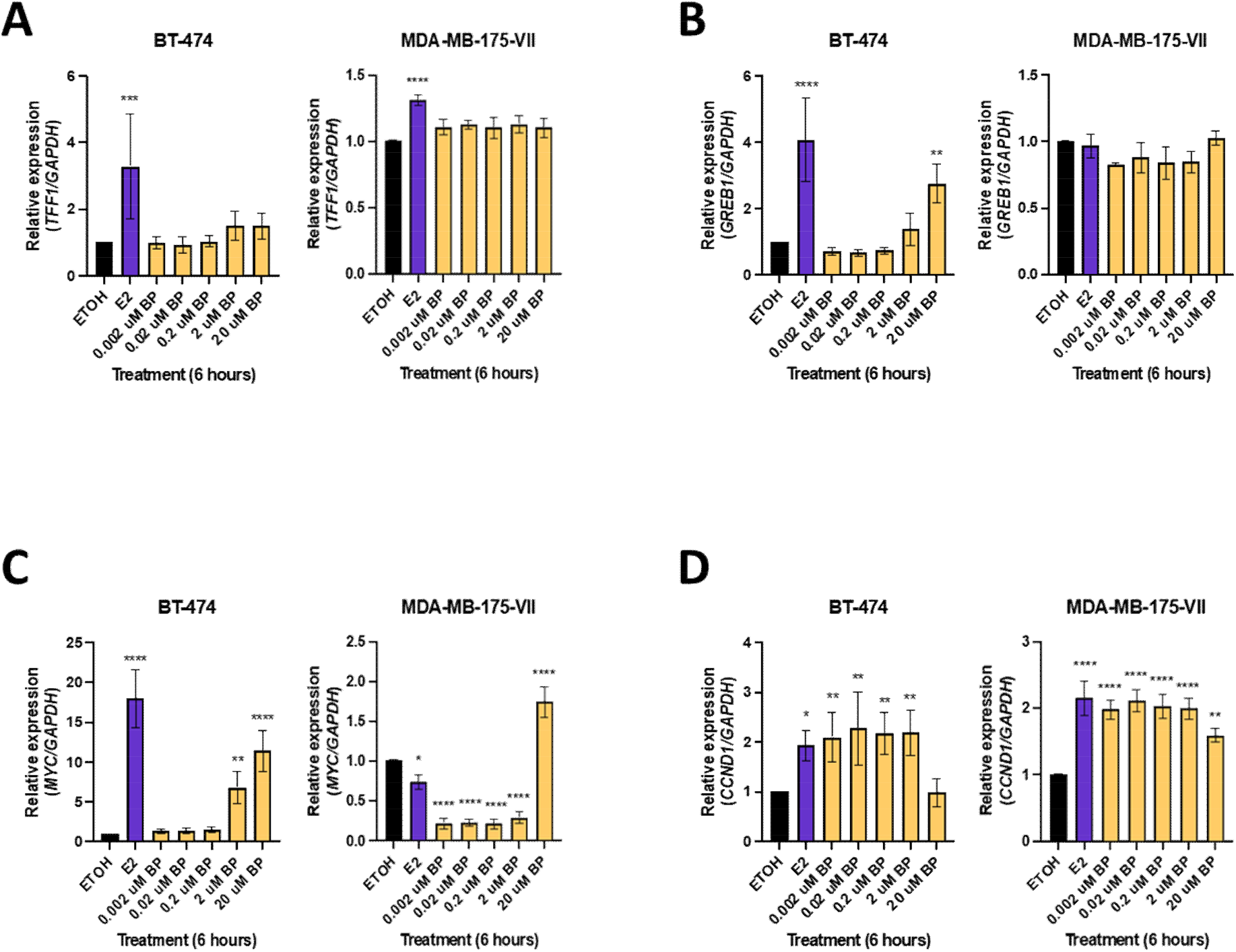
BP-mediated ER target gene expression in BT-474 and MDA-MB-175-VII cells. BT-474 and MDA-MB-175-VII luminal B breast cancer cells were treated with biologically relevant doses of BP for 6 hours. Quantification of **(A) *TFF1*, (B) *GREB1*, (C) *MYC*, and (D) *CCND1*** gene expression following BP exposure. *PGR* was not expressed in the MDA-MB-175-VII cell line. EtOH was used as a negative control, and E2 (10 nM) served as a positive control. GAPDH was used as a housekeeping gene. n=3, *p<0.05, **p<0.01, one-way ANOVA.

**Supplemental Figure 5.**
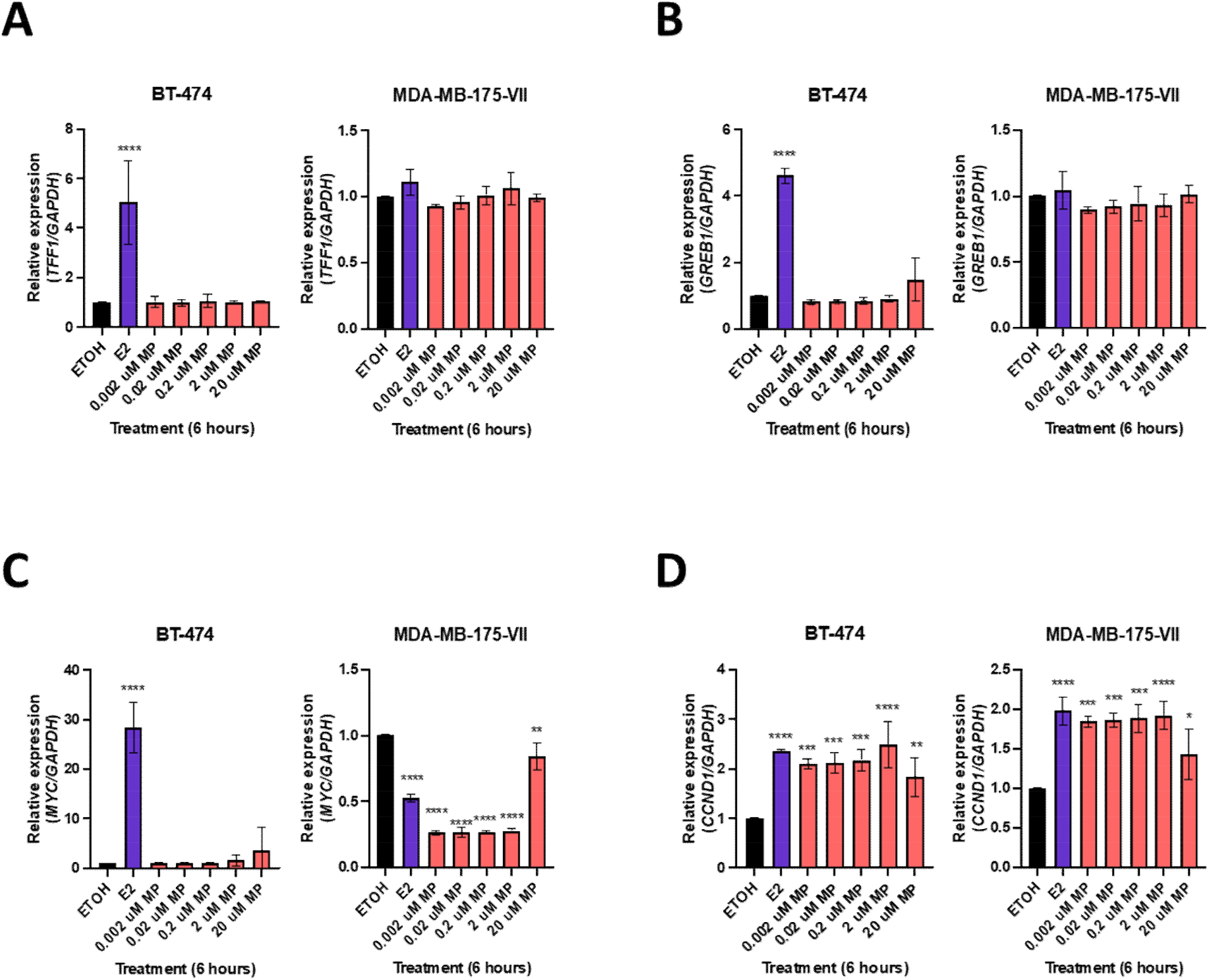
MP-mediated ER target gene expression in BT-474 and MDA-MB-175-VII cells. BT-474 and MDA-MB-175-VII luminal B breast cancer cells were treated with the indicated doses of MP for 6 hours. Quantification of **(A)** *TFF1*, **(B)** *GREB1, MYC*, and **(D)** *CCND1* gene expression following BP exposure. *PGR* was not expressed in the MDA-MB-175-VII cell line. EtOH was used as a negative control, and E2 (10 nM) served as a positive control. GAPDH was used as a housekeeping gene. n=3, *p<0.05, **p<0.01, one-way ANOVA.

**Supplemental Figure 6.**
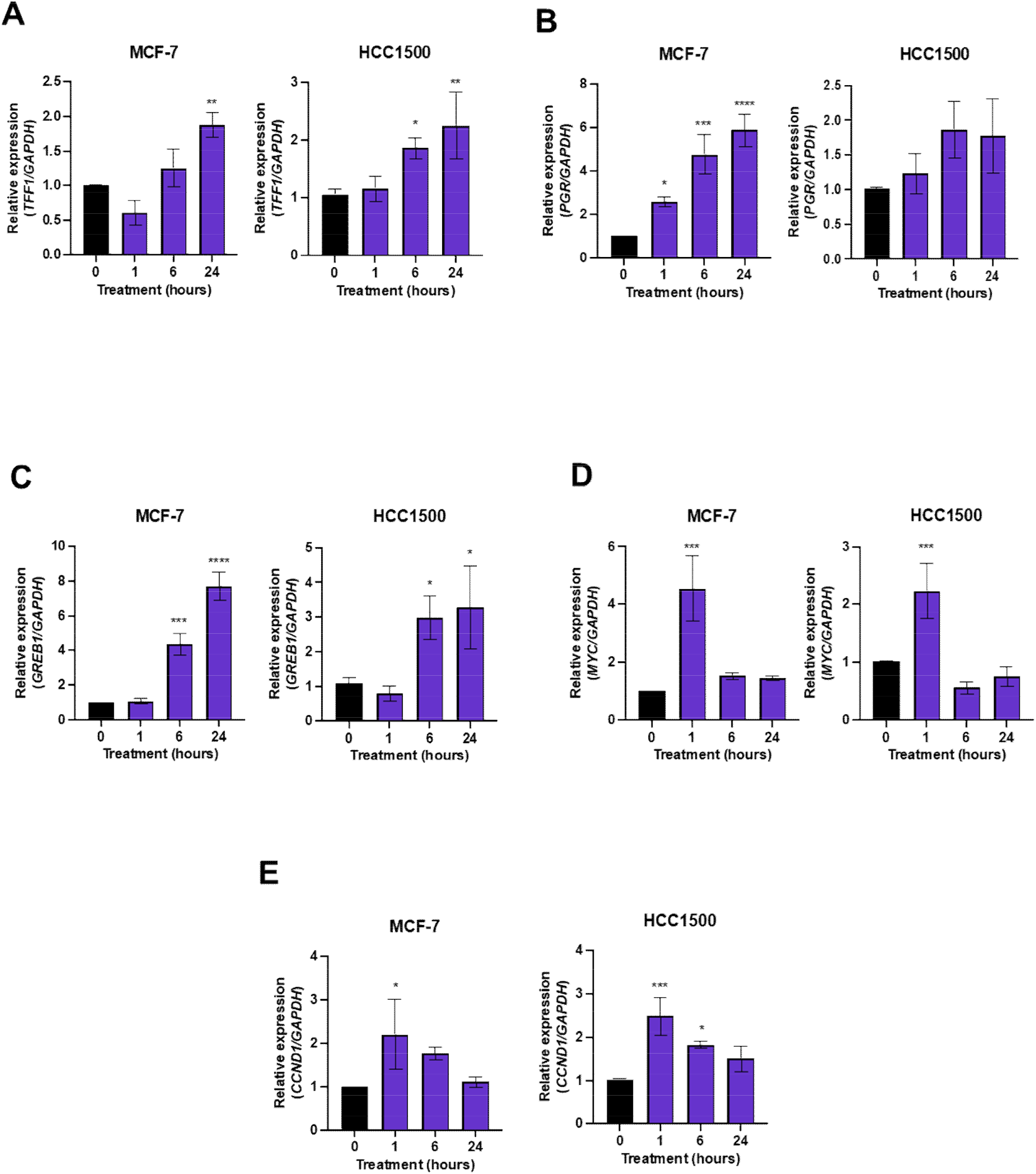
E2-mediated regulation of ER target gene expression is time dependent. MCF-7 and HCC1500 luminal A breast cancer cells were treated with E2 (10 nM) for 1, 6, or 24 hours. Relative gene expression of **(A)** *TFF1*, **(B)** *PGR*, **(C)** *GREB1*, **(D)** *MYC*, and **(E)** *CCND1* was assessed by real-time qualitative polymerase chain reaction. EtOH was used as a negative control. GAPDH was used as a housekeeping gene. n=3, *p<0.05, **p<0.01, ***p<0.001, one-way ANOVA.

**Supplemental Figure 7.**
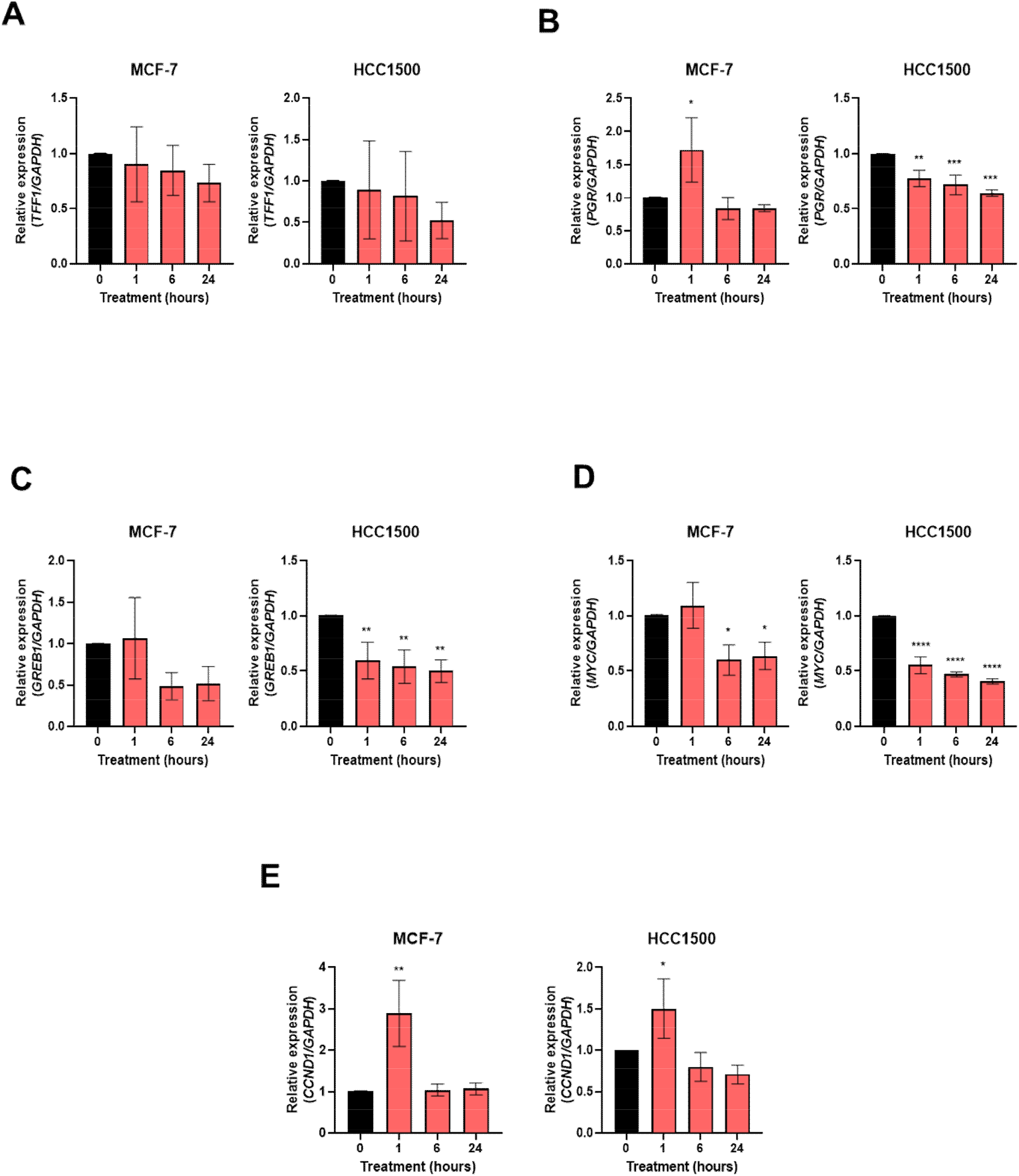
MP-mediated regulation of ER target gene expression is time dependent. MCF-7 and HCC1500 luminal A breast cancer cells were treated with MP (2 µM) for 1, 6, or 24 hours. Relative gene expression of **(A)** *TFF1*, **(B)** *PGR*, **(C)** *GREB1*, **(D)** *MYC*, and **(E)** *CCND1* was assessed by real-time qualitative polymerase chain reaction. EtOH was used as a negative control, and E2 (10 nM) served as a positive control. GAPDH was used as a housekeeping gene. n=3, *p<0.05, **p<0.01, ***p<0.001, one-way ANOVA.

**Supplemental Figure 8.**
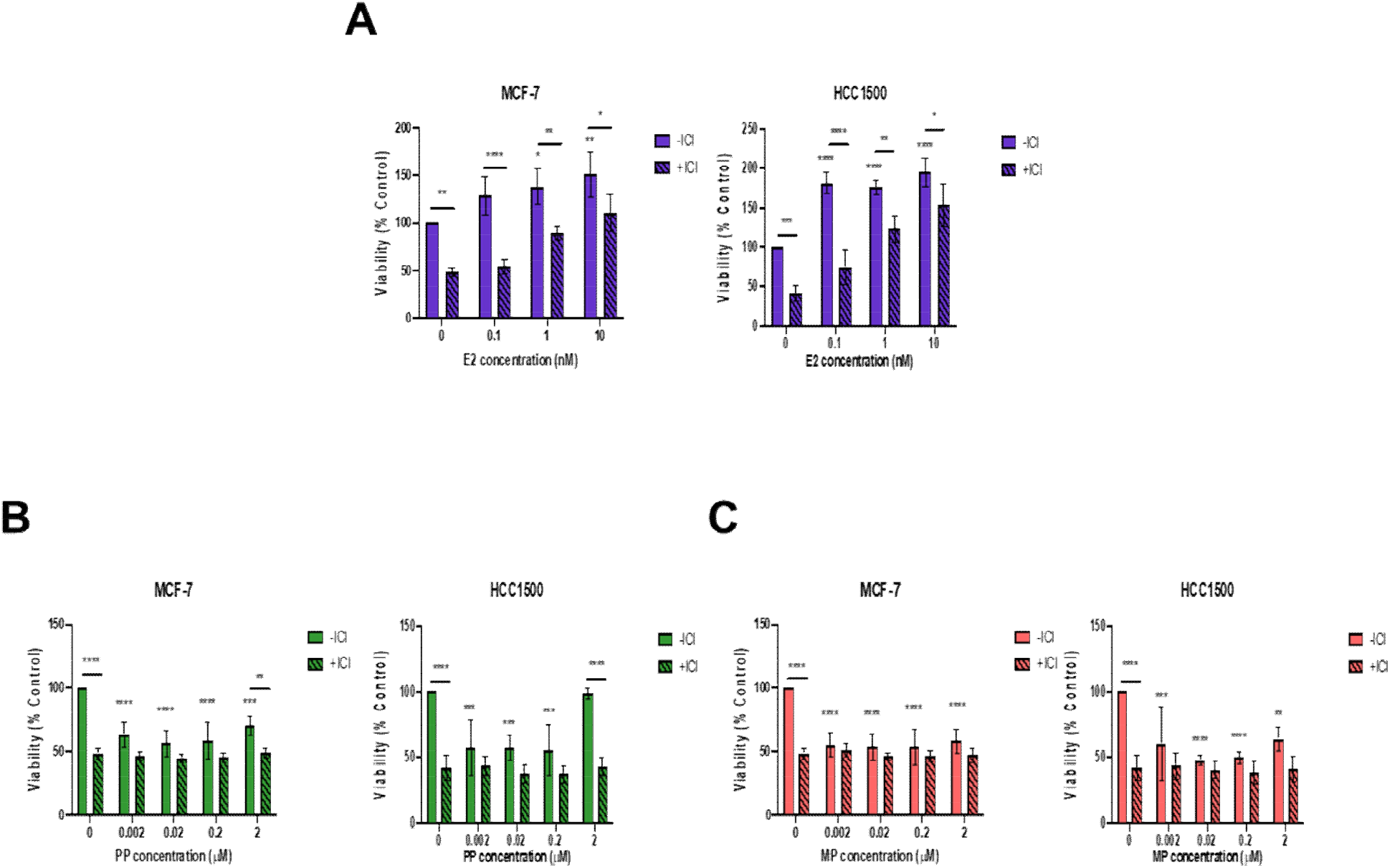
Effect of PP or MP exposure on cell viability is not ER-dependent. MCF-7 and HCC1500 luminal A breast cancer cells were co-treated with ER-antagonist, ICI 182, 780 (1 nM), and **(A)** E2, **(B)** PP, or **(C)** MP at the indicated doses for 7 days. EtOH was used as a negative control. n=4, *p<0.05, **p<0.01, ***p<0.001, ****p<0.0001, two-way ANOVA.

**Supplemental Figure 9.**
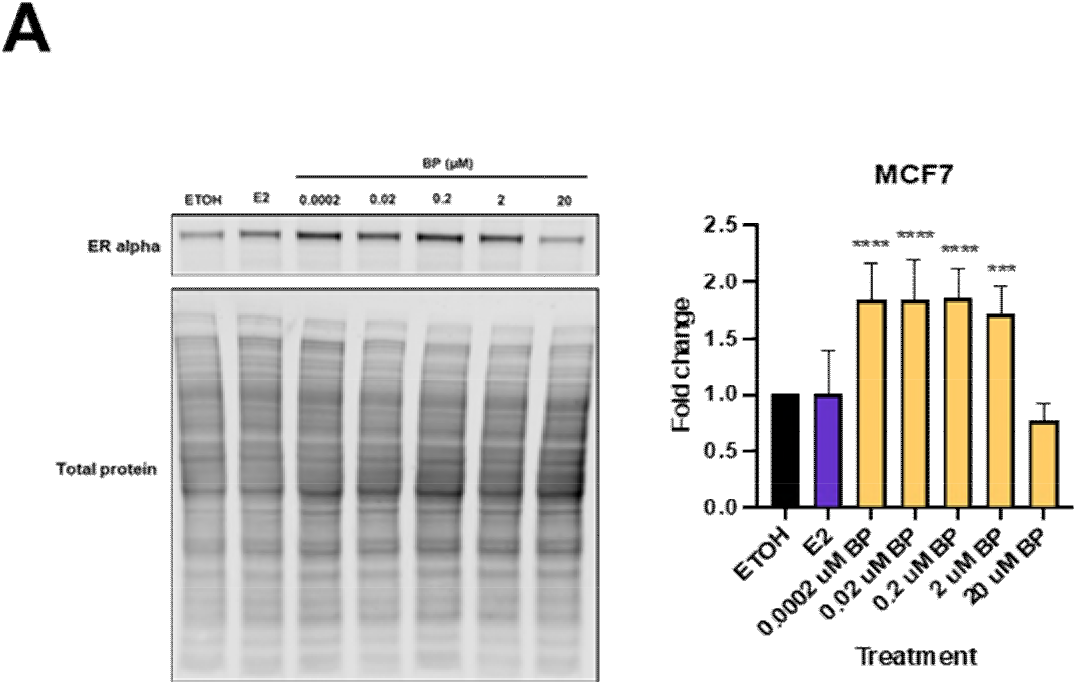
BP-mediated effects on ER protein expression in MCF-7 cells. MCF-7 luminal A breast cancer cells were treated with the indicated doses of BP for 24 hours. **(A)** Representative western blot image, and quantified bar graph displaying BP-mediated effect on ERα protein expression in the MCF-7 cell line. EtOH was used as a negative control, and E2 (10 nM) served as a positive control. n=6, **p<0.01, ***p<0.001, one-way ANOVA.

